# Investigating the role of behaviour in the genetic risk for schizophrenia

**DOI:** 10.1101/611079

**Authors:** Jessye Maxwell, Adam Socrates, Kylie P. Glanville, Marta Di Forti, Robin M. Murray, Evangelos Vassos, Paul F. O’Reilly

## Abstract

The notion that behaviour may be on a causal path from genetics to psychiatric disorders, such as schizophrenia, highlights a potential for practical interventions. Motivated by this, we test the association between schizophrenia (SCZ) polygenic risk scores (PRS) and 420 behavioural traits (personality, psychological, lifestyle, nutritional) in a psychiatrically healthy sub-cohort of the UK Biobank. Higher schizophrenia PRS was associated with a range of traits, including lower verbal-numerical reasoning (*P* = 6×10^−61^), higher nervous feelings (*P* = 2×10^−51^) and higher self-reported risk-taking (*P* = 2×10^−41^). We follow-up the risk-taking association, hypothesising that the association may be due to a genetic propensity for risk-taking leading to greater migration, urbanicity or drug-taking – reported environmental risk factors for schizophrenia, and all positively associated with risk-taking in these data. However, schizophrenia PRS was also associated with traits, such as tea drinking (*P* = 2×10^−34^), that are highly unlikely to be on a causal path to schizophrenia. We depict four causal relationships that may in theory underlie such PRS-trait associations and illustrate ways of testing for each. For example, we contrast PRS-trait trends in the healthy sub-cohort to the corresponding trait values of medicated and non-medicated individuals diagnosed with schizophrenia, allowing some differentiation of mediation-by-behaviour, disease-onset effects and treatment effects. However, dedicated follow-up studies and new methods are required to fully disentangle these relationships. Thus, while we urge caution in interpretation of simple PRS cross-trait associations, we propose that well-designed PRS analyses can contribute to identifying behaviours on the causal path from genetics to disease.

---

It is now clear that pleiotropy is a ubiquitous feature of the human genome^1–4^. Polygenic risk scores (PRS), which correspond to a weak proxy of an individual’s genetic propensity to a trait or disease, can be used to estimate the extent of genetic overlap between phenotypes via predicting one phenotype from the PRS of another^5^. PRS have been applied, as have bivariate GCTA-GREML^6^ and LD score regression^2^, to estimate shared genetic aetiology across hundreds of phenotypes^2,7–9^.

Exploiting the UK Biobank, we identify associations between PRS for schizophrenia and a range of behavioural traits, including cognitive, personality and nutritional measures (**Fig. 1**). We follow-up a schizophrenia PRS vs risk-taking association, observing phenotypic associations between risk-taking and substance abuse and within-UK migration patterns, which may indicate a path from genetics to schizophrenia via risk-taking behaviour. Whether such a path reflects a common route to schizophrenia requires further investigation but highlights the potential of such cross-trait analyses. However, there are also highly significant associations with traits such as cooked vegetable intake and tea drinking, which we use to illustrate how different causal explanations must be considered (**Fig. 2**) before concluding simple mediation-by-behaviour mechanisms from PRS-trait associations.

**Figure 1.**
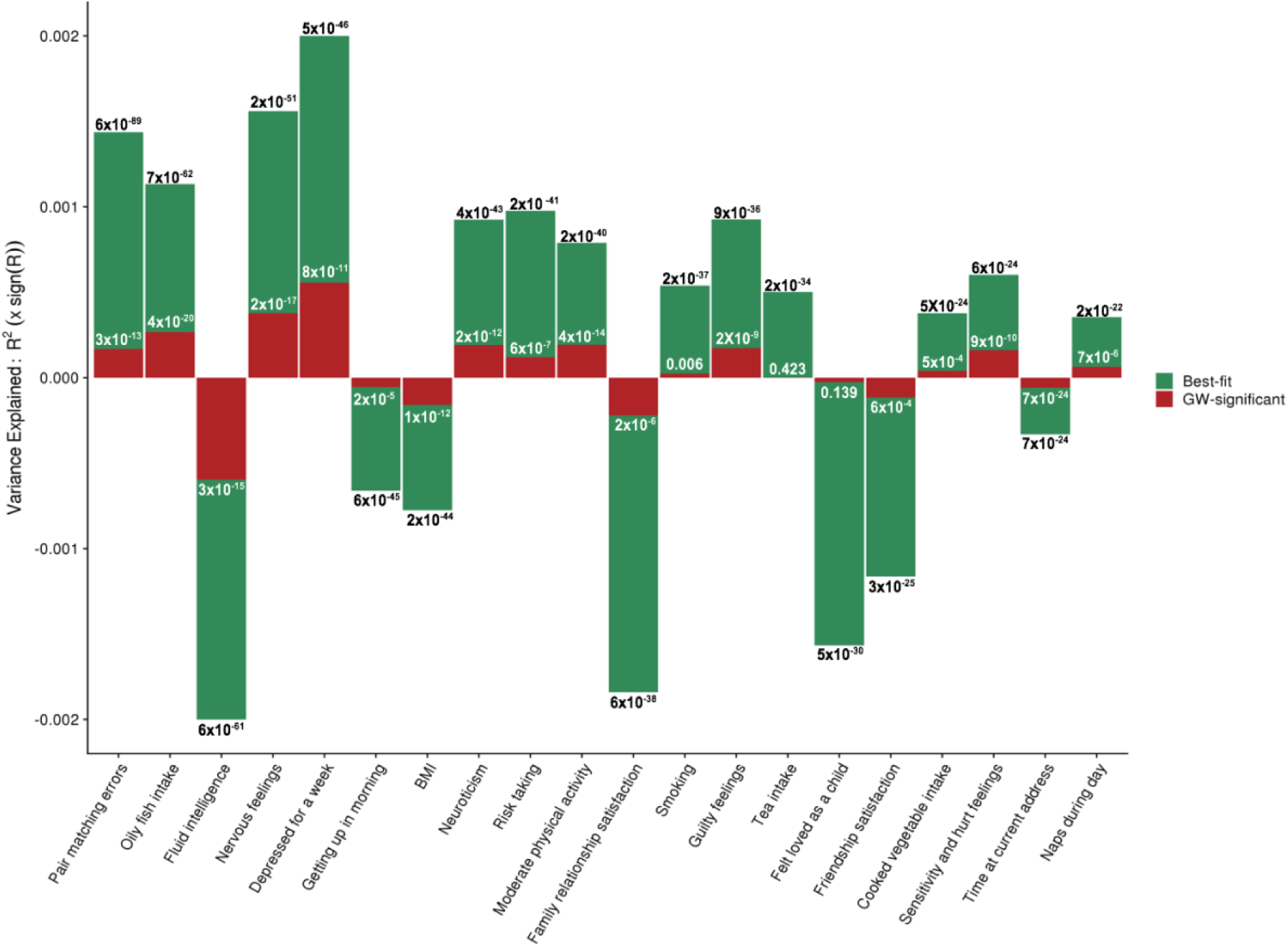
Bar plot showing 20 of the most significant associations (Y-axis shows target trait variance explained, upwards for positive and downwards for negative associations; with *P*-values of association shown on bars) between the best-fit SCZ PRS (green) and the corresponding target traits (X-axis), as well as the association results when using the genome-wide significant (GW-significant) SCZ PRS instead (based on the 108 sentinel SCZ SNPs). See **Supp. Table 1** for details and to inspect the selection of these 20 results among the top results.

**Figure 2.**
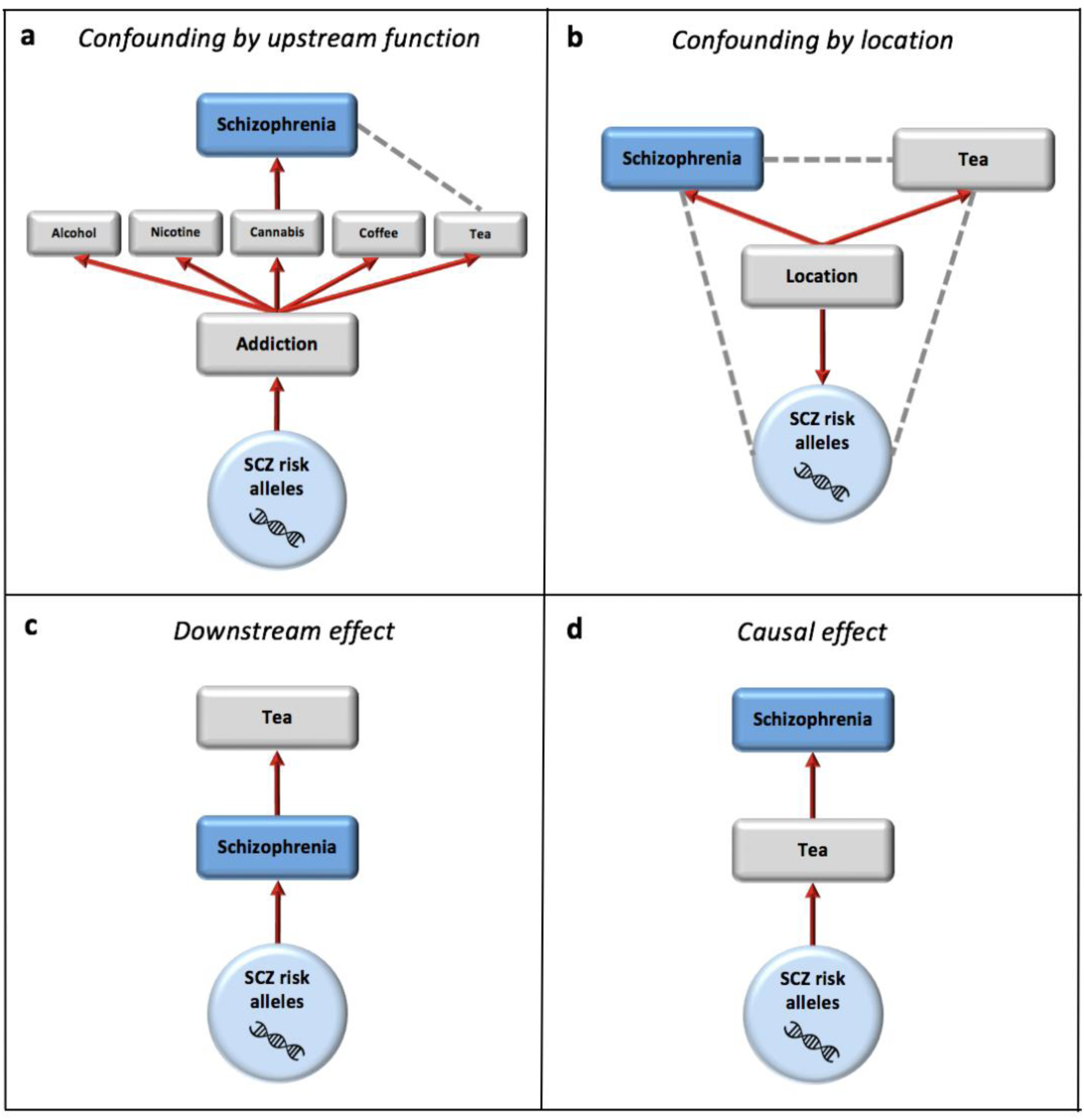
Four possible causal explanations for the SCZ-PRS vs tea drinking association. Red arrows reflect causal paths and grey dotted lines depict confounded associations induced by the causal relationships. SCZ ‘risk alleles’ refer to those included in the PRS, thus comprising genuine SCZ risk alleles (/risk haplotypes) and those with no effect on SCZ. **a.** Some fraction of SCZ risk alleles are addiction risk alleles, which influence addiction to multiple substances. The causal relationship between cannabis and schizophrenia assumed here induces the association between SCZ-PRS and tea consumption, confounded by function (addiction here, used for illustration only); **b.** SCZ risk allele frequencies vary with location (due to genetic drift etc) and so too do environmental factors influencing tea consumption and SCZ risk, inducing correlations between all three; **c.** SCZ risk alleles increase risk for SCZ. Schizophrenia increases tea consumption; **d.** SCZ risk alleles increase tea consumption. Tea consumption increases risk for SCZ.

A key feature of our study design is that the UK Biobank sample size allows us to contrast PRS vs behaviour trends in the unaffected population with the same behaviours in 985 individuals with schizophrenia, in the same cohort. With PRS an incomplete measure of genetic liability, we investigate whether the associations between increasing polygenic risk for schizophrenia and behavioural traits in the unaffected population are predictive of the trait values in medicated and non-medicated schizophrenia cases, who we expect to have higher genetic liability on average (**Fig. 3**). This provides insights into the potential effects of schizophrenia genetic risk, disease onset, and medication, on behaviour across the liability spectrum. Therefore, analyses of this type could act as a complement to other approaches, such as Mendelian Randomisation, for investigating causality in trait-disease associations^10^.

**Figure 3.**
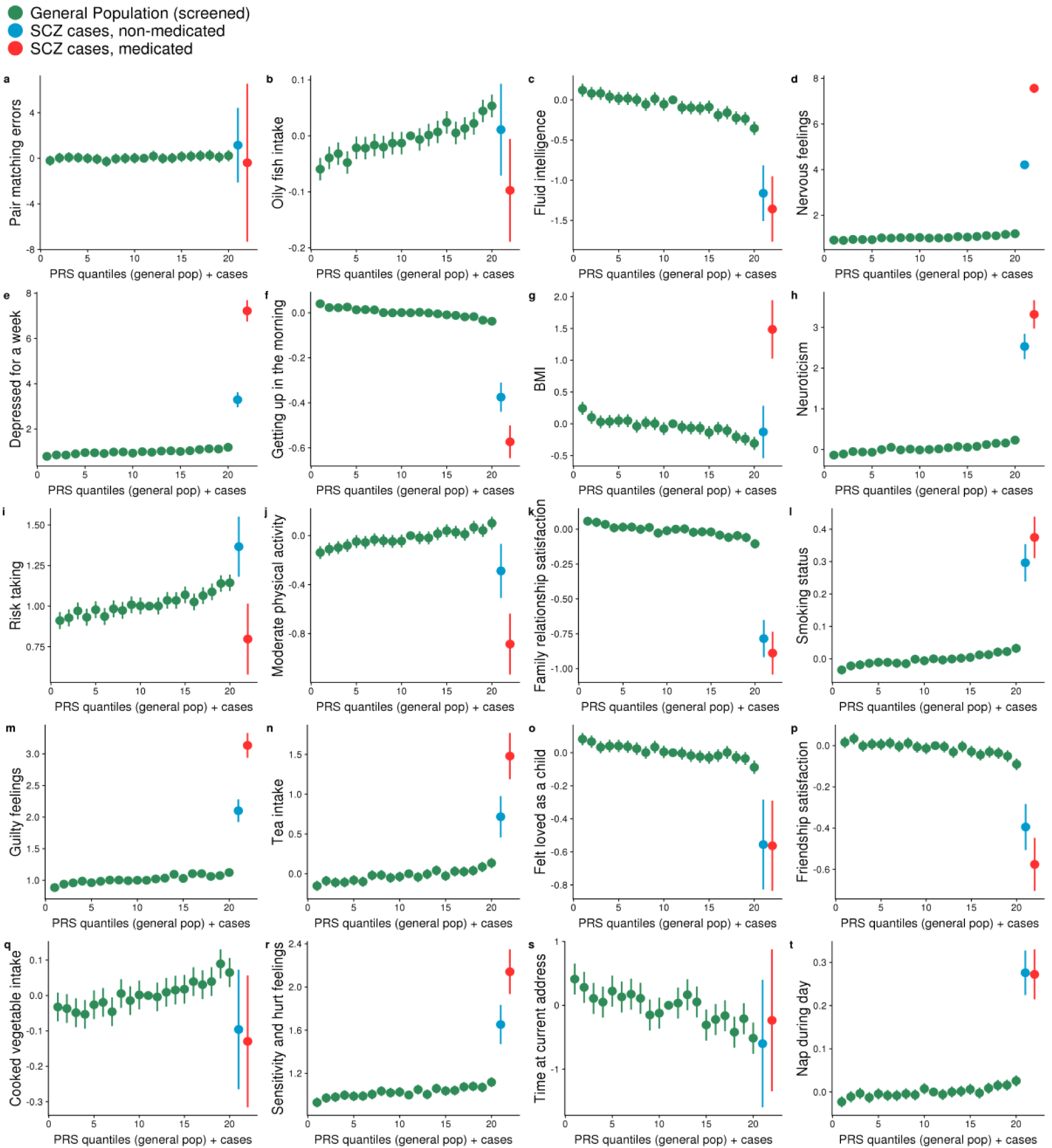
PRS-by-trait quantile plots (in green) indicating the trends of association between the best-fit SCZ PRS and the target behavioural traits among 20 of the most significant associations, matching those of **Fig. 1**. The quantile plot is ordered from low to high genetic risk for schizophrenia, according to PRS in the target (UK Biobank) data, with each green point representing the average target trait value of a 5% quantile of the sample. Non-medicated (blue) and medicated (red), at baseline, diagnosed individuals are appended to the right end of each plot, reflecting the expected higher genetic burden of diagnosed individuals compared to unaffected individuals (see Main Text). Vertical lines represent 95% confidence intervals; these appear absent for some traits with a large range and are larger in the two categories of cases due to their smaller sample sizes.

## RESULTS

We performed PRS analyses (see **Methods**) using PRSice^11^, with the schizophrenia (SCZ) GWAS summary statistics from the Psychiatric Genomics Consortium (PGC) as base data^12^ and the UK Biobank data on 307,823 unaffected individuals as target data (see **Supp. Table 1** for traits and sample sizes). Associations between the most predictive SCZ PRS and the 420 behavioural traits from the UK Biobank were tested using linear and logistic regressions in PRSice, controlling for age, sex, Townsend deprivation index and the first 15 principal components to control for population stratification. We set a conservative significance threshold of *P* < 1×10^−10^ based on testing the most predictive PRS (according to SNP inclusion at different *P*-value thresholds; see **Methods**) across 420 target behavioural traits.

SCZ PRS showed significant (*P* < 1×10^−10^) associations in 101 of the 420 behavioural traits in the unaffected sub-cohort of 307,823 individuals. To verify the reliability of these polygenic associations, we repeated the analyses using PRS based only on the 108 sentinel genome-wide significant schizophrenia variants^12^. 77 of the 101 associated traits were nominally significant (*P* < 0.05) based on the PRS comprising only genome-wide significant variants. The results of all 101 significant associations are displayed in **Supp. Table 1**.

**Fig. 1** illustrates 20 of the most significant associations, with highly related traits omitted (see **Supp. Table 1**), showing both the results based on the best-fit PRS (green) and the PRS calculated from only the 108 genome-wide significant SCZ SNPs (red). Here the higher explanatory power of best-fit PRS is apparent, while the consistency with the GW-significant PRS is reassuring. Most of the associations are in a direction that reflects ‘negative trait’ outcomes with higher genetic risk for SCZ; for example, positive correlations with nervous (*P* = 1×10^−51^) and guilty (8×10^−36^) feelings, and negative correlations with fluid intelligence (*P* = 5×10^−61^) and friendship satisfaction (*P* = 3×10^−25^).

### A pathway to schizophrenia via the genetics of risk-taking

Focusing on one of the associations that may be most amenable to intervention, that with self-reported risk-taking (response to question “Would you describe yourself as someone who takes risks?’), we hypothesise that part of the aetiology of schizophrenia may derive from a genetic propensity for risk-taking, resulting in greater exposure to drug-taking, migration or urbanicity – reported risk factors for schizophrenia^13–16^. Testing these pathways explicitly is underpowered here due to the relatively low power of a UK Biobank risk-taking PRS and the small number of schizophrenia cases, so instead we tested the phenotypic associations and those with the SCZ PRS. Migration patterns were characterised in three ways: Euclidean distance moved between birth and current residence, birth and current residence population density (and their difference), and time spent at current residence (see **Methods**). We also tested associations with several ‘control traits’, such as ‘leg pain on walking’, ‘breast fed’ and ‘month attended baseline assessment centre’ (see **Methods**) for comparison, since epidemiological testing on such large sample sizes are vulnerable to producing highly significant associations that are the consequence of small biases or statistical artefacts.

Testing in unaffected individuals, self-reported risk-taking is positively associated with lifetime distance moved (*P* = 3×10^−123^; *r*^*2*^ = 0.001), change in population density (*P* = 7×10^−37^; *r*^*2*^ = 0.0004) and negatively associated with time spent in current residence (*P* < 1×10^−300^; *r*^*2*^ = 0.007). Risk-taking is also positively associated with self-reported substance abuse (*P* = 8×10^−74^; *r*^*2*^ = 0.008). While these results are consistent with the proposed pathway, we caution against their strong interpretation because of their small explanatory power, their attenuation when also controlling for Townsend deprivation index and education level, the sensitivity of small *P*-values to stochastic variation^17^, and because most of the associations with the control traits were also significant (see **Supp. Table 2** for all results).

Next, we tested for the association between schizophrenia PRS and these traits in the unaffected sub-cohort (see **Supp. Table 3**). With the exception of left hand grip strength, BP device ID and population density of birth place, all the nominally significant results related to the migration variables and substance use and all the non-significant results corresponded to the control traits (threshold of *P* < 0.05 for this sub-analysis, though we caution discrete interpretation of *P*-values in general^18^). Schizophrenia PRS was positively associated with population density of current residence (*P* = 3×10^−11^), self-reported substance abuse (4×10^−9^), ever smoked cannabis (7×10^−8^) and change in population density from birth to current residence (*P* = 6×10^−3^), and negatively associated with time at current residence (*P* = 1×10^−21^) and distance travelled (5×10^−4^). However, each association is attenuated when Townsend deprivation index and education are controlled for, especially lifetime change in population density. This may reflect a complex network of causal relationships between the genetics of risk-taking, education, socio-economic status and migration and drug-taking patterns, which could generate cross-generational effects such as the association observed between SCZ PRS and birthplace population density. Devoted follow-up studies will be required to further unravel the proposed risk-taking – schizophrenia pathway.

### Potential confounding-by-function in PRS associations

While many of the associations in **Fig. 1** have been implicated in schizophrenia aetiology previously or are consistent with the notion that liability to schizophrenia can cause subtle deficits^19^, there are also highly significant unexpected associations between the best-fit SCZ PRS and tea-intake (*P* = 1×10^−34^) and cooked vegetable intake (*P* = 4×10^−24^). While the association between the GW-significant PRS and tea intake is reassuringly non-significant (*P* = 0.42), it remains a concern for the many similar analyses being conducted in the field^7–9^ that there is such a significant association between the best-fit PRS and a trait that is expected to play no causal role in schizophrenia. We exploit the engaging and memorable nature of this example to highlight the potential confounding in such analyses, and to illustrate what we consider to be four main causal explanations for this association (**Fig. 2**).

While **Fig. 2a** corresponds to what is usually termed ‘horizontal pleiotropy’, we think it useful to illustrate one (hypothetical) example of how this may manifest, since the idea that a genetic variant that affects schizophrenia risk also happens to affect tea drinking seems unlikely without a mediating shared function (a hypothesised addiction pathway used here for illustrative purpose). The relationship of **Fig. 2a** could be tested by controlling for cannabis smoking, but relatively few unaffected individuals in the UK Biobank answered the questions on cannabis smoking (N = 32,125 had tried cannabis), such that the original association between SCZ-PRS and tea drinking is no longer significant in that sub-cohort. However, we find positive associations phenotypically between tea drinking and cannabis smoking (*P* = 6×10^−4^; maximum frequency of cannabis smoking), and also substance abuse (*P* = 0.05), although alcohol consumption (*P* = 3×10^−106^) and coffee consumption (*P* < 1×10^−308^) are negatively correlated with tea drinking. Also, while the best-fit SCZ PRS is positively associated with current smoking status (*P* = 2×10^−37^) and ever-taken-cannabis (*P* = 7×10^−17^), consistent with the literature^20,21^, it is negatively associated with beer intake (*P* = 2×10^−11^) and coffee intake (*P* = 3×10^−12^). Thus, any link with propensity to addiction is not straightforward and unlikely to be explained by the simple relationship in **Fig. 2a**. Nevertheless, the consequences of confounding-by-function could be widespread: a range of functions other than addiction, such as cognition, anxiety, impulsivity or cellular processes such as synaptic pruning, that may be on the causal pathway to schizophrenia, could generate associations between genetic risk for schizophrenia and numerous non-causal factors in the general population.

The potential for confounding by location (**Fig. 2b**) in polygenic score analyses has been highlighted in several recent publications^5,22,23^. We repeated the analyses extending the number of principal components (PCs) adjusted for from 15 to 40 PCs and observed little change in results (**Supp. Fig. 1**). While confounding by location is, thus, perhaps unlikely to be the main explanation for the association with tea drinking, the large sample size here ensures that even subtle confounding effects – not well-captured by top PCs – could be responsible for highly significant associations, and so without a thorough investigation of potential confounding by population structure this possibility cannot be ruled out.

While individuals with schizophrenia may consume more tea (**Fig. 2c**) – due to spending more time indoors, for example – our analyses were performed in the unaffected sub-cohort and so the observed association should not be due to disease onset. However, we discuss the possibility of this, due to misclassification or the spectrum-like manifestation of the disease in the general population, in the next section.

We believe that tea being a causal risk factor for schizophrenia (**Fig. 2d**) is the least likely of these potential explanations. Bidirectional Mendelian Randomization^24^ and the Latent Causal Variable approach^25^ can be applied to distinguish between causal effects in either direction (**Fig. 2c** and **Fig. 2d**), but distinguishing these from confounding-by-function (**Fig. 2a**) is likely to prove extremely challenging without highly rich or prospective data.

Each of the models in **Fig. 2** are severe simplifications of reality, which likely involves simultaneous bidirectional and feedback effects, but are presented here to highlight some key different causal relationships consistent with observed PRS trait associations.

### Effects of disease onset and medication

The presence of individuals with schizophrenia in the UK Biobank allows us to contrast the PRS-by-trait trends observed in the unaffected general population with values of the corresponding traits in diagnosed individuals. There are 1151 individuals in the full UK Biobank data with an ICD-10 diagnosis (from Hospital Episodes Statistics) of, or self-reported, schizophrenia. 728 of these individuals reported presently taking antipsychotic treatment during the baseline verbal interview, which is also when most traits tested here were measured; however, many traits were measured as part of the Mental Health Questionnaire performed approximately 8 years after baseline interviews and so there will be greater misclassification of treatment status for these traits and correspondingly smaller differences (**Supp. Table 1**). **Fig. 3** displays the PRS-by-trait trends in the unaffected general population as quantile plots (in green), with the trait values for non-medicated (blue) and medicated (red) SCZ cases appended to the right end of the X-axes. Depicting the cases as having the highest genetic liability is appropriate because, on average, cases are expected to have a higher genetic liability than unaffected individuals, and the low predictive power of the PRS ensures that even individuals in the top 5% quantile of the unaffected sub-cohort will only have a moderately elevated schizophrenia risk on average.

The plots in **Fig. 3** correspond to the traits of **Fig. 1** and are shown in the same order of association significance (notice the different Y-axis scales when comparing the trends).

In **Fig. 3d** we observe higher levels of nervous feelings in unaffected individuals with higher genetic risk for SCZ, even higher levels in non-medicated SCZ cases, and the highest levels of nervous feelings in medicated SCZ cases. While BMI reduces with increasing SCZ PRS (**Fig. 3g**), medicated cases have markedly higher BMI (*P* = 1×10^−5^; see **Supplementary Table 1**), consistent with the known side-effect of anti-psychotic medication of increased BMI. **Fig. 3j** may point to some behavioural explanation for these BMI associations, since higher SCZ PRS is associated with higher physical activity, while medicated individuals show the lowest levels of activity. While there is increased self-reported risk-taking with higher SCZ PRS (**Fig. 3i**), individuals on medication have lower levels of risk-taking than those not on medication (*P* = 3×10^−3^): interestingly, dopaminergic drugs have been found to increase risky behaviours in several studies on Parkinson’s disease patients^26,27^ and so these results, pertaining to anti-psychotic dopaminergic-inhibitors may reflect the reverse effect.

In 13 of the 20 results shown in **Fig. 3**, the trait values in SCZ cases are in the direction expected according to the PRS-by-trait trend. Misclassification of SCZ cases as being unaffected, or the pre-clinical or spectrum-like manifestation of the disorder in the general population, could theoretically have generated the results relating to these 13 traits. For example, if misclassification of SCZ cases as unaffected is more common among individuals with higher SCZ PRS, then a downstream effect on eg. tea consumption (**Fig. 2c**) may have generated the observed PRS-by-trait positive trend. However, if such misclassification, or pre-clinical, effects exist, then they cannot explain all the results since almost half (7 of 20) show discordant results between the general and affected sub-cohorts. **Supp. Figs. 2-5** display similar plots for all PRS-by-trait association results with *P* < 1×10^−10^, and **Supp. Table 1** provides results of testing the difference between trait values in SCZ cases compared to those of the top quantile in the unaffected general population.

The widespread significant and substantial differences observed in the behavioural traits between schizophrenia cases and the unaffected individuals (**Fig.3, Supp. Table 1, Supp. Figs. 2-5**): (1) supports the recorded diagnosis of schizophrenia (ie. that these individuals behave differently from the general population), and (2) stresses the need to perform PRS cross-trait analyses that are stratified by case/medication status.

## DISCUSSION

This study exploited GWAS data from the Psychiatric Genomics Consortium (PGC) and genetic and phenotype data from the UK Biobank, to examine the associations between the common genetic liability for schizophrenia, estimated by polygenic risk scores, and a range of behavioural traits. Approximately one quarter of the associations, which were observed in unaffected individuals past the typical age of schizophrenia diagnosis (> 40 years), were significant at a stringent threshold. Most of these show that increased genetic risk of schizophrenia is associated with negative trait outcomes, such as lower friendship and family satisfaction and greater feelings of guilt and anxiety, reflecting known social cognition deficits in diagnosed and high-risk individuals^28,29^. Thus, the broad findings appear to support the notion that common genetic risk for schizophrenia manifests as a continuum across the population, with symptoms common to diagnosed schizophrenia cases observed in high risk unaffected individuals to a greater extent than low risk unaffected individuals. This finding may seem trivial but need not have been the case, and, if true, is more consistent with these symptoms being on the causal path to schizophrenia than being downstream effects of disease onset, although confounding-by-function (**Fig. 2a**) is also possible.

While there is justified concern about generating false-positive associations when exploiting polygenic scores that include a large number of null variants, we observed high consistency between the results of best-fit PRS and PRS based only on genome-wide significant SNPs (**Fig. 1**), with best-fit PRS having markedly higher explanatory power. Atypical differences between the results of the two types of PRS could highlight potentially confounded associations, as in the case of tea drinking. However, a significant best-fit PRS and non-significant GW-significant PRS may also reflect a peripheral but causal effect, whereby a trait that is only weakly causal of schizophrenia is influenced by variants that are correspondingly of very small effect and thus not among the GW-significant SNPs. Thus, as well as increasing explanatory power, the results from best-fit PRS could provide aetiological insights if evaluated carefully.

The unprecedented size of the UK Biobank as a deeply genotyped-phenotyped data set allowed us to perform a novel type of analytical comparison. The data were of sufficient size to both conduct a well-powered investigation of PRS-trait relationships at high resolution and to contrast those relationships with corresponding trait values of a sufficient number of individuals with schizophrenia (**Fig. 3**), all within the same cohort. This within-study comparison provides a unique insight into comparative general population, disease onset and medication effects. The contrast between the trends of association in the unaffected and affected sub-cohorts for a fraction of the traits demonstrated that all results cannot be only a consequence of disease misclassification or pre-clinical effects, although these effects are likely to contribute to an extent to the observed associations. We hope that our analyses will motivate the collection of samples with sufficiently large numbers of both unaffected and affected individuals, consistently assayed and ideally prospectively collected, so that effects across the disease liability spectrum can be explored. The potential for gaining causal insights from analyses such as these, which include no cryptic assumptions, should make them a useful counterpart to other approaches for causal inference, such as Mendelian Randomisation^10,24^. In our future work we plan to develop a formal method of causal inference based on this design.

However, while there is information to be gleaned from such cross-trait PRS analyses, the results must be interpreted carefully. The trait variance explained by the SCZ PRS has a maximum of only 0.04% (**Supp. Table 1**) and for most traits tested is much lower, which means that (1) at best we have some initial insights into the role of behaviour in the genetic risk for schizophrenia, and (2) the generation of misleading results due to subtle bias or confounding cannot be entirely ruled out. Although many of the leading associations, and their directions, are consistent with the literature on schizophrenia aetiology, we highlight the need for more thorough investigation before strong interpretation of any specific association. We illustrate four simple associations (**Fig. 2**) that could plausibly generate any of the observed PRS-trait associations. Focusing on tea drinking as an example may seem farcical, and was in fact the topic of a popular blogpost highlighting the potential problems of PRS associations (with tea drinking a *hypothetical* example)^30^, but is intended to clearly demonstrate the fallacy of assuming that highly significant PRS-trait associations only expose factors on the causal pathway, potentially amenable to intervention. For this reason, a large-scale systematic study of this type must be considered as exploratory, providing only broad aetiological insights and a screen of potentially interesting links between traits and disease requiring further investigation.

Exemplifying the kind of follow-up analyses that can be performed based on the results of such exploratory studies, we further investigated the association of SCZ-PRS and self-reported risk-taking by calculating lifetime within-UK migration patterns and combining those with population density data from the UK National Office of Statistics. These analyses indicated that risk-taking genetics may be a sub-component of the genetic aetiology of schizophrenia due to mediation by migration and/or drug-taking^13–16^. However, confirmation of this and an assessment of the contribution of such a relationship to overall schizophrenia prevalence requires dedicated follow-up studies. It is interesting to note that genetic-environment correlations of this type effectively inflate both pedigree-based and population-based heritability estimates, since they are regarded as a genetic contribution to the phenotype even if the causal agent is environmental.

This study represents arguably the most systematic interrogation of the link between genetic risk for schizophrenia and behaviour to date. We have generated a catalogue of 420 schizophrenia PRS – behavioural trait associations, 101 exceeding a stringent multiple testing threshold. While the large-scale nature of the analyses makes them necessarily exploratory, we have demonstrated several ways of gaining greater insights than those derived from the main associations. We hope that these analytical strategies, and the array of PRS-trait associations generated here, will act as a useful starting point for follow-up investigations to expose behavioural traits on the causal path from genetics to schizophrenia, which may provide targets of early intervention.

## METHODS

### Base and target data

The base (or discovery) data for this study were the genome-wide association study (GWAS) results from the Psychiatric Genomics Consortium (PGC) for schizophrenia^12^. These schizophrenia GWAS results are derived from a sample of up to 36,989 cases and 113,075 controls of European and East Asian ancestry, which yielded 108 independent loci harbouring genome-wide significant associations.

In the present study we analysed target data on up to 307,823 participants (52% females), aged 38 – 73 years (mean = 56.85, S.D. = 8.06), involved in the UK Biobank baseline assessment (http://www.ukbiobank.ac.uk)^31^ and Mental Health Questionnaire. UK Biobank is a health resource for researchers that aims to improve the prevention, diagnosis and treatment of a range of illnesses. The recruitment process was coordinated around 22 centres in the UK (between 2007 and 2010)^32^. Individuals within travelling distance of these centres were identified using NHS patient registers (response rate = 5.47%)^33^. Invitations were sent using a stratified approach to ensure demographic parameters were in concordance with the general population. All participants provided written consent and the current study was ethically approved by the UK Biobank Ethics and Governance Council (REC reference 11/NW/0382; UK Biobank application reference 18177).

The traits analysed in the target data were “behavioural traits”, liberally defined as any trait that has a substantial behavioural component and included traits across personality, mood, nutrition, physical activity and psychological feelings. Altogether 420 such behavioural traits were analysed, although many of these traits were closely related to each other and thus there is a strong correlation structure among the traits. Details on 101 of these traits are contained in **Supp. Table 1**.

### Base and target data: Quality control and exclusions

Blood samples from 488,366 UK Biobank participants were genotyped using the UK BiLEVE array or the UK Biobank axiom array. Further details on the genotyping and quality control (QC) can be found on the UK Biobank website (http://www.ukbiobank.ac.uk/scientists-3/genetic-data/). In the current study, SNPs were removed if they had missingness < 0.02 and MAF < 0.01. Samples were removed from the dataset if they had missingness < 0.01. A subset of European ancestry inferred individuals were defined using 4-means clustering applied to the first two principal components of the genotype data. One of each pair of related individuals were removed using a relatedness criterion *KING* coefficient < 0.088. Exclusions based on heterozygosity and missingness were implemented according to UK Biobank recommendations (http://biobank.ctsu.ox.ac.uk/showcase/label.cgi?id=100314). Samples were removed if they were discordant for sex. SNPs deviating from Hardy-Weinburg equilibrium (HWE) were removed at a threshold of *P* < 10^−8^. This QC process resulted in a data set of 560,173 SNPs and 386,192 samples available for analysis.

For our polygenic risk analyses of the unaffected general population, we removed all individuals with an ICD-10 diagnosis of major depressive disorder, bipolar disorder, schizophrenia, and all individuals on antipsychotic medication. There were 307,823 individuals remaining for the analyses of SCZ PRS in the unaffected sub-cohort. The analyses that performed comparisons between the unaffected sample and individuals diagnosed with SCZ exploited data on 985 individuals in the full UK Biobank data with an ICD-10 hospital diagnosis of schizophrenia and their antipsychotic medication status. 438 individuals of these individuals reported antipsychotic treatment during the verbal interview.

### Primary polygenic risk score analyses

Summary statistics from the SCZ GWAS were downloaded from the PGC website (https://www.med.unc.edu/pgc/results-and-downloads). Polygenic risk scores (PRS) were generated using PRSice v2 software (www.prsice.info)^11^. PRSice calculates individual risk scores by calculating the sum of disease-associated alleles, weighted by the log odds ratio estimated in the discovery GWAS. SNPs were clumped to minimise their linkage disequilibrium (LD) using an r2 ≥ 0.1 threshold in sliding windows of 250 kb. PRS on SCZ were generated for individuals in the UK Biobank cohort based on the PGC discovery GWAS summary statistics, and then used to predict target phenotypes recorded the UK Biobank. To find the most predictive model for each of the target traits, PRS were calculated at 2001 thresholds between *P* = 5×10^−8^ and 1 at increments of 0.0005. Under a Bonferroni correction for multiple testing that conservatively assumes independence between PRS thresholds and among the base and target traits, we use a significance threshold of 0.05 / (2001*420) = 6×10^−8^ for the most predictive PRS in our primary analyses, based on the 2001 PRS thresholds and 420 target traits tested. For further stringency, given the risk of detecting significant associations that are a result of subtle confounding in such large sample sizes, we only declare significance at *P* < 1×10^−10^. Note that the selection of the most predictive PRS means that the trait variance explained (R^2^) by the best-fit PRS will be inflated here but since the aim of our study is to gain broad aetiological insights rather than for direct clinical utility, which requires far higher predictive power, then this should not affect any of our conclusions. To obtain unbiased estimates of trait variance explained then out-of-sample prediction or cross-validation should be performed^5^. Details of the 101 phenotypes that show an association with the most predictive PRS at *P* < 1×10^−10^ in the target data are provided in **Supp. Table 1**. Associations were examined in regression models for the binary (logistic) and quantitative (linear) target traits, adjusting for age, sex, Townsend deprivation index and the first 15 principal components of the genotype data to control for population stratification, with additional adjustments described in the text for relevant analyses. The principal components were provided as part of the UK Biobank genotype data release.

### Migration and substance abuse analyses

The UK Biobank data included birth and current residence (at baseline) location variables, under our application, defined using the British National Grid referencing system, corresponding to northerly and easterly positions, with a reference point close to the Isles of Scilly. We used these locations to characterise migration patterns of UK Biobank individuals within the UK. First, the Euclidean distance travelled between birth and current residence was calculated using the northerly/easterly co-ordinates of each; the distance is calculated to the nearest kilometre to prevent identification (mean distance travelled = 84.9km, median = 15.0km). Next, population density measures were derived by matching the co-ordinates of each participant to population density information held on national databases relating to each local authority district. Population density statistics and boundary data for local authority districts were downloaded from the office of national statistics (https://www.ons.gov.uk), based on the 2011 UK Census data. The R packages sp and rgdal were then used to retrieve the appropriate population density metrics for each participant according to their location co-ordinates. Thus, the population density measures for birth place (mean = 2376 per km^2^, median = 1897 per km^2^), current residence (mean = 2059 per km^2^, median = 1415 per km^2^), and their difference (mean = −316.7, median = 0), were derived for each participant and, again, corresponded to the local postcode area. Time spent at current residence was available directly as a UK Biobank variable (mean = 17.8 years, median = 16 years). Substance abuse, defined as those with an ICD-10 hospital diagnosis or self-reported substance abuse (total UK Biobank sample: cases = 4033, controls = 498631), was also available.

Self-reported risk-taking was tested for association with the migration phenotypes using linear regression and with substance abuse using logistic regression. The control phenotypes that were used in this follow-up analyses, also tested via linear and logistic regression, were: breast fed (binary: cases = 277671, controls = 106156); birth weight known (binary: cases = 277076, controls = 224726); birth weight (mean = 3.12kg, median = 3.32kg); left hand grip (mean = 29.55, median = 28); leg pain on walking (cases = 39759, control = 130710); blood pressure device ID and month attended assessment centre. Each of these phenotypes were then tested for an association with the SCZ PRS. All of the analyses were adjusted for age, sex, Townsend deprivation index and educational attainment, as described in the main text and corresponding table captions.

