## Supplementary material for "Investigating the role of behaviour in the genetic risk for schizophrenia"

**Supplementary Table 1**: Target traits associated with SCZ PRS at *P* < $1x10^{-10}$ in the unaffected sub-cohort of the UK Biobank.

| Trait | Direction of effect | Best-fit $R^2$ | Best-fit PRS *P*-value | GW-Sig PRS *P*-value | *N* | Unaffected vs cases difference *P*-value |
|---|---|---|---|---|---|---|
| **Number of incorrect matches in round 2 (*Pair matching errors*)** | **+** | **0.0013** | **$6x10^{-89}$** | **$3x10^{-13}$** | **307395** | **0.72** |
| Number of fluid intelligence questions attempted within time limit | - | 0.0036 | $3x10^{-81}$ | $1x10^{-27}$ | 99466 | 0.03 |
| Seen doctor GP for nerves anxiety tension or depression | + | 0.0016 | $6x10^{-77}$ | $2x10^{-15}$ | 305607 | $2x10^{-08}$ |
| Plays computer games | - | 0.001 | $3x10^{-68}$ | $3x10^{-08}$ | 307479 | $4x10^{-06}$ |
| Seen a psychiatrist for nerves anxiety tension or depression | + | 0.0024 | $5x10^{-65}$ | $1x10^{-16}$ | 306579 | $4x10^{-235}$ |
| Time to complete round 2 | + | 0.0009 | $5x10^{-62}$ | $1x10^{-16}$ | 307395 | $1x10^{-03}$ |
| **Oily fish intake** | **+** | **0.0009** | **$7x10^{-62}$** | **$4x10^{-20}$** | **306254** | **$5x10^{-04}$** |
| **Fluid intelligence score⁺** | **-** | **0.0026** | **$6x10^{-61}$** | **$3x10^{-15}$** | **99466** | $2x10^{-17}$ |
| Reaction time | + | 0.0008 | $2x10^{-60}$ | $2x10^{-07}$ | 305797 | $4x10^{-24}$ |
| **Nervous feelings** | **+** | **0.0012** | **$2x10^{-51}$** | **$2x10^{-17}$** | **300070** | **$2x10^{-67}$** |
| Frequency of tenseness restlessness in last 2 weeks | + | 0.0007 | $3x10^{-50}$ | $2x10^{-09}$ | 296554 | $5x10^{-51}$ |
| Time to complete round 1 | + | 0.0007 | $5x10^{-49}$ | $9x10^{-11}$ | 307395 | $6x10^{-03}$ |
| **Ever depressed for a whole week** | **+** | **0.0027** | **$5x10^{-46}$** | **$8x10^{-11}$** | **99725** | **$4x10^{-30}$** |
| Duration to first press of snap button in each round 1 | + | 0.0006 | $6x10^{-46}$ | $4x10^{-08}$ | 305103 | $4x10^{-16}$ |
| Duration to first press of snap button in each round 2 | + | 0.0006 | $2x10^{-45}$ | $4x10^{-05}$ | 305279 | $1x10^{-07}$ |
| Ever unenthusiastic disinterested for a whole week | + | 0.0029 | $4x10^{-45}$ | $8x10^{-10}$ | 97754 | $1x10^{-40}$ |
| **Getting up in morning** | **-** | **0.0006** | **$6x10^{-45}$** | **$2x10^{-05}$** | **307056** | **$3x10^{-37}$** |
| Number of incorrect matches in round 1 | + | 0.0006 | $2x10^{-44}$ | $1x10^{-07}$ | 307395 | $2x10^{-09}$ |
| **BMI** | **-** | **0.0006** | **$2x10^{-44}$** | **$1x10^{-12}$** | **306919** | **$5x10^{-18}$** |
| **Neuroticism score (*Neuroticism*)** | **+** | **0.0007** | **$4x10^{-43}$** | **$2x10^{-12}$** | **250454** | **$5x10^{-64}$** |
| Reason for reducing amount of alcohol drunk | - | 0.0016 | $3x10^{-42}$ | $4x10^{-03}$ | 112644 | $2x10^{-06}$ |
| **Self-reported risk taking (*Risk taking)*** | **+** | **0.0009** | **$2x10^{-41}$** | **$6x10^{-07}$** | **297101** | **< $1x10^{-308}$** |
| Number of symbol digit matches attempted | - | 0.002 | $6x10^{-41}$ | $4x10^{-06}$ | 68773 | $1x10^{-05}$ |

| | | | | | | |
|---|---|---|---|---|---|---|
| **Number of days week of moderate physical activity 10 minutes (*Moderate physical activity*)** | **+** | **0.0006** | **$2x10^{-40}$** | **$4x10^{-14}$** | **293844** | **$2x10^{-09}$** |
| **Family relationship satisfaction** | **+** | **0.0016** | **$6x10^{-38}$** | **$2x10^{-06}$** | **100952** | **$1x10^{-22}$** |
| **Current smoking status (*Smoking*)** | **+** | **0.0005** | **$2x10^{-37}$** | **$6x10^{-03}$** | **306505** | **$3x10^{-09}$** |
| Worrier anxious feelings | + | 0.0007 | $7x10^{-36}$ | $3x10^{-13}$ | 299507 | $1x10^{-19}$ |
| Duration to first press of snap button in each round 0 5 | + | 0.0005 | $7x10^{-36}$ | $6x10^{-06}$ | 304982 | $6x10^{-23}$ |
| **Guilty feelings** | **+** | **0.0008** | **$9x10^{-36}$** | **$2x10^{-09}$** | **299852** | **$5x10^{-25}$** |
| **Tea intake** | **+** | **0.0005** | **$2x10^{-34}$** | **0.42** | **297475** | **$1x10^{-03}$** |
| Frequency of depressed mood in last 2 weeks | + | 0.0005 | $9x10^{-33}$ | $7x10^{-10}$ | 295142 | $2x10^{-54}$ |
| Time to answer | + | 0.0007 | $2x10^{-32}$ | $7x10^{-09}$ | 101682 | 0.03 |
| Non oily fish intake | + | 0.0004 | $8x10^{-32}$ | $3x10^{-12}$ | 306525 | $6x10^{-07}$ |
| Duration to first press of snap button in each round 11 | + | 0.0004 | $1x10^{-30}$ | 0.02 | 304995 | $8x10^{-27}$ |
| **Felt loved as a child** | **-** | **0.0015** | **$5x10^{-30}$** | **0.14** | **83605** | **$5x10^{-04}$** |
| Frequency of unenthusiasm disinterest in last 2 weeks | + | 0.0004 | $1x10^{-29}$ | $3x10^{-07}$ | 298216 | $9x10^{-51}$ |
| Past tobacco smoking | - | 0.0004 | $3x10^{-29}$ | $2x10^{-04}$ | 288806 | $3x10^{-01}$ |
| Duration to first press of snap button in each round | + | 0.0009 | $2x10^{-28}$ | $9x10^{-04}$ | 125518 | $3x10^{-05}$ |
| Frequency of tiredness lethargy in last 2 weeks | + | 0.0004 | $4x10^{-27}$ | $4x10^{-06}$ | 298429 | $2x10^{-42}$ |
| Number of correct matches in round 2 | - | 0.0004 | $1x10^{-25}$ | $1x10^{-03}$ | 307395 | $8x10^{-23}$ |
| **Friendships satisfaction** | **+** | **0.001** | **$3x10^{-25}$** | **$6x10^{-04}$** | **100886** | **$7x10^{-12}$** |
| Time to complete round | + | 0.0015 | $4x10^{-25}$ | $4x10^{-04}$ | 66954 | $1x10^{-08}$ |
| PM initial answer | - | 0.001 | $1x10^{-24}$ | $3x10^{-06}$ | 101664 | $6x10^{-12}$ |
| **Cooked vegetable intake** | **+** | **0.0003** | **$5x10^{-24}$** | **$5x10^{-04}$** | **298869** | **0.08** |
| **Sensitivity and hurt feelings** | **+** | **0.0004** | **$6x10^{-24}$** | **$9x10^{-10}$** | **298692** | **$5x10^{-12}$** |
| **Length of time at current address** | **-** | **0.0003** | **$7x10^{-24}$** | **$3x10^{-06}$** | **300865** | **$5x10^{-04}$** |
| Ever had prolonged feelings of sadness or depression | + | 0.0016 | $1x10^{-23}$ | $4x10^{-03}$ | 83668 | $3x10^{-24}$ |
| **Naps during day** | **+** | **0.0003** | **$2x10^{-22}$** | **$7x10^{-06}$** | **307469** | $5x10^{-34}$ |
| Duration to first press of snap button in each round 0 4 | + | 0.0007 | $2x10^{-22}$ | $8x10^{-04}$ | 125055 | $1x10^{-07}$ |
| Repeated disturbing thoughts of stressful experience in past month | + | 0.0011 | $2x10^{-22}$ | 0.03 | 83808 | 0.16 |
| Time spent watching television TV | - | 0.0003 | $4x10^{-22}$ | 0.60 | 290951 | 0.08 |
| Duration screen displayed | + | 0.0009 | $7x10^{-22}$ | $2x10^{-06}$ | 101682 | $5x10^{-09}$ |
| Bipolar and major depression status | + | 0.0012 | $8x10^{-22}$ | $2x10^{-07}$ | 73573 | $2x10^{-25}$ |
| Number of correct matches in round 1 | - | 0.0003 | $1x10^{-21}$ | 0.01 | 307395 | $1x10^{-21}$ |

| | | | | | | |
|---|---|---|---|---|---|---|
| Avoided activities or situations because of previous stressful experience in past month | + | 0.0011 | $1x10^{-21}$ | 0.07 | 83805 | $4x10^{-09}$ |
| Number of days week of vigorous physical activity 10 minutes | + | 0.0003 | $2x10^{-21}$ | $1x10^{-05}$ | 293799 | $2x10^{-08}$ |
| Pork intake | - | 0.0003 | $2x10^{-21}$ | $4x10^{-03}$ | 306068 | 1 |
| Number of vehicles in household | - | 0.0003 | $2x10^{-21}$ | 0.02 | 305805 | $1x10^{-166}$ |
| Ever smoked | + | 0.0004 | $3x10^{-21}$ | $5x10^{-04}$ | 306540 | $6x10^{-10}$ |
| Happiness | + | 0.0008 | $1x10^{-20}$ | $8x10^{-06}$ | 101565 | $2x10^{-18}$ |
| Duration to first press of snap button in each round 3 | + | 0.0007 | $2x10^{-20}$ | $9x10^{-03}$ | 125546 | $3x10^{-04}$ |
| Ever felt worried tense or anxious for most of a month or longer | + | 0.0018 | $4x10^{-20}$ | $3x10^{-03}$ | 80410 | $1x10^{-12}$ |
| Age first had sexual intercourse | - | 0.0003 | $5x10^{-20}$ | $5x10^{-04}$ | 269875 | 0.90 |
| Sleep duration | + | 0.0003 | $1x10^{-19}$ | $3x10^{-04}$ | 306190 | $6x10^{-39}$ |
| Felt very upset when reminded of stressful experience in past month | + | 0.0009 | $6x10^{-19}$ | 0.46 | 83794 | $4x10^{-08}$ |
| Duration to first press of snap button in each round 2 | + | 0.0006 | $1x10^{-18}$ | $2x10^{-03}$ | 126174 | $2x10^{-06}$ |
| Ever taken cannabis | + | 0.0008 | $7x10^{-18}$ | $6x10^{-04}$ | 83787 | 0.02 |
| Overall health rating | + | 0.0002 | $3x10^{-17}$ | 0.30 | 306588 | $2x10^{-93}$ |
| Worry too long after embarrassment | + | 0.0003 | $4x10^{-17}$ | 0.03 | 295015 | $8x10^{-19}$ |
| General happiness | + | 0.0008 | $5x10^{-17}$ | 0.07 | 83653 | $1x10^{-08}$ |
| Ever suffered mental distress preventing usual activities | + | 0.0015 | $1x10^{-16}$ | 0.08 | 82792 | $2x10^{-55}$ |
| Loneliness isolation | + | 0.0004 | $7x10^{-16}$ | 0.02 | 303135 | $7x10^{-59}$ |
| Qualifications university | + | 0.0003 | $1x10^{-15}$ | 0.34 | 307812 | $3x10^{-24}$ |
| Qualifications GCSEs | - | 0.0003 | $3x10^{-15}$ | 0.26 | 307812 | 0.03 |
| Current tobacco smoking | + | 0.0002 | $3x10^{-15}$ | 0.17 | 307490 | $3x10^{-35}$ |
| Belittlement by partner or ex partner as an adult | + | 0.0007 | $5x10^{-15}$ | 0.37 | 83687 | $1x10^{-05}$ |
| Number of incorrect matches in round | + | 0.0009 | $9x10^{-15}$ | 0.06 | 68804 | 0.81 |
| Physical violence by partner or ex partner as an adult | + | 0.0007 | $2x10^{-14}$ | 0.58 | 83707 | 0.01 |
| Never eat eggs dairy wheat sugar wheat | - | 0.0009 | $7x10^{-47}$ | $7x10^{-47}$ | 307812 | $4x10^{-07}$ |
| Felt hated by family member as a child | + | 0.0007 | $6x10^{-14}$ | 0.27 | 83737 | $8x10^{-06}$ |
| Duration to first press of snap button in each round 1 | + | 0.0004 | $8x10^{-14}$ | 0.07 | 126495 | $5x10^{-24}$ |
| Number of pregnancy terminations | + | 0.0011 | $2x10^{-13}$ | $2x10^{-03}$ | 48343 | $2x10^{-03}$ |
| Number of attempts | + | 0.0005 | $2x10^{-13}$ | 0.01 | 101682 | $1x10^{-03}$ |
| Ever had prolonged loss of interest in normal activities | + | 0.0006 | $4x10^{-13}$ | $1x10^{-04}$ | 83678 | $1x10^{-27}$ |
| Length of mobile phone use | - | 0.0002 | $7x10^{-13}$ | 0.38 | 303951 | $6x10^{-42}$ |

| | | | | | | |
|---|---|---|---|---|---|---|
| Ever thought that life not worth living | + | 0.0006 | $2x10^{-12}$ | 0.18 | 83477 | $1x10^{-17}$ |
| Mood swings | + | 0.0002 | $3x10^{-12}$ | $4x10^{-04}$ | 300051 | $2x10^{-43}$ |
| Coffee intake | - | 0.0002 | $3x10^{-12}$ | $1x10^{-05}$ | 285407 | $5x10^{-09}$ |
| Recent feelings of nervousness or anxiety | + | 0.0006 | $3x10^{-12}$ | $4x10^{-05}$ | 83610 | $6x10^{-08}$ |
| Loud music exposure frequency | + | 0.0005 | $3x10^{-12}$ | 0.02 | 100541 | $5x10^{-05}$ |
| FI3 word interpolation | - | 0.0005 | $4x10^{-12}$ | $4x10^{-03}$ | 98680 | $3x10^{-05}$ |
| Ever worried more than most people would in similar situation | + | 0.0012 | $5x10^{-12}$ | 0.01 | 72425 | $4x10^{-18}$ |
| Ever sought or received professional help for mental distress | + | 0.0009 | $7x10^{-12}$ | 0.01 | 83656 | $< 1x10^{-308}$ |
| Ever contemplated self harm | + | 0.0005 | $1x10^{-11}$ | 0.11 | 83704 | $4x10^{-13}$ |
| Number of unsuccessful stop smoking attempts | + | 0.0007 | $1x10^{-11}$ | 0.04 | 68272 | 0.41 |
| Beef intake | - | 0.0001 | $2x10^{-11}$ | 0.12 | 306645 | $2x10^{-05}$ |
| Morning evening person chronotype | + | 0.0002 | $3x10^{-11}$ | 0.39 | 274298 | $3x10^{-11}$ |
| Age at first episode of depression | - | 0.0012 | $5x10^{-11}$ | 0.28 | 32234 | $5x10^{-06}$ |
| Ever highly irritable argumentative for 2 days | + | 0.0007 | $7x10^{-11}$ | 0.13 | 99716 | $4x10^{-11}$ |
| Victim of physically violent crime | + | 0.0005 | $9x10^{-11}$ | 0.08 | 83793 | $4x10^{-17}$ |

GW-Sig PRS = PRS comprising only genome-wide significant ($P<5x10^{-8}$). Top non-overlapping and distinct traits in bold selected for **Fig. 1** of the Main Text (*names given in Figure 1 provided in parentheses if different*). [+] Fluid intelligence assessed by a simple 13 question verbal-numerical reasoning test. Final column shows the result (*P*-value) of a difference-in-means t-test between the target trait value among individuals diagnosed with schizophrenia and with unaffected individuals.

**Supplementary Table 2**: Associations between self-reported risk-taking and measures of migration, substance use and control traits

| | Effect (β): adjusted for age and sex | $R^2$: adjusted for age and sex | *P*-value: adjusted for age and sex | *P*-value: adjusted for age, sex and TDEP | *P*-value: adjusted for age, sex, TDEP and EDU |
|---|---|---|---|---|---|
| Ever smoked cannabis | 0.23 | 0.11 | < $10^{-308}$ | < $10^{-308}$ | < $10^{-308}$ |
| Time at current residence | -2.23 | 0.00664 | < $10^{-308}$ | < $10^{-308}$ | < $10^{-308}$ |
| Left hand grip strength | 0.651 | 0.000627 | 1.32xl0$^{-143}$ | 1.23xlO$^{-200}$ | 9.95x10$^{-181}$ |
| Population density: current residence | 204 | 0.00149 | 3.19x10$^{-136}$ | 4.94xl0$^{-50}$ | 7.28xl0$^{-24}$ |
| Distance travelled: birth to current | 10800 | 0.00131 | 2.61xl0$^{-123}$ | 1.14x10$^{-133}$ | 2.38xl0$^{-67}$ |
| Self-reported substance abuse | 1.85$^{+}$ | 0.00771$^{\alpha}$ | 7.94xl0$^{-74}$ | 5.52xl0$^{-55}$ | 3.93xl0$^{-62}$ |
| Breast fed | 1.16$^{+}$ | 0.00115$^{\alpha}$ | 4.67xl0$^{-67}$ | 2.39xl0$^{-68}$ | 5.95xl0$^{-51}$ |
| Leg pain on walking | 1.23$^{+}$ | 0.00229$^{\alpha}$ | 2.55xl0$^{-56}$ | 3.85xl0$^{-38}$ | 1.52xl0$^{-52}$ |
| Population density difference: birth to current | 127 | 0.000389 | 7.13xl0$^{-37}$ | 2.31x10$^{-09}$ | 0.0252 |
| Population density: birth place | 76.8 | 0.000198 | 1.50x10$^{-19}$ | 1.67x10$^{-10}$ | 2.44x10$^{-10}$ |
| Birth weight | 0.0254 | 0.000271 | 1.39x 10$^{-17}$ | 4.53xl0$^{-20}$ | 1.45x10$^{-16}$ |
| Birth weight known | 0.96$^{+}$ | 9.45xl0$^{-05\alpha}$ | 2.80x10$^{-09}$ | 0.000935 | 4.39xl0$^{-05}$ |
| BP device ID | 156000 | 2.50x10$^{-06}$ | 0.277 | 0.267 | 0.331 |
| Month attended baseline assessment | -0.00327 | 1.87xl0$^{-07}$ | 0.766 | 0.778 | 0.86 |

+ Odds ratio for binary outcome variables; α Pseudo $R^2$ for binary outcome variables; TDEP = Townsend deprivation index; EDU = Educational attainment

**Supplementary Table 3**: Summary of PRS results corresponding to the most predictive PRS for SCZ tested against the target behavioural and control traits

| | Effect (β): adjusted for age and sex | $R^2$: adjusted for age and sex | *P*-value: adjusted for age and sex | *P*-value: adjusted for age, sex and TDEP | *P*-value: adjusted for age, sex, TDEP and EDU |
|---|---|---|---|---|---|
| Time at current residence | -738.61 | 0.00027 | 1.13x10$^{-21}$ | 2.32x10$^{-21}$ | 9.07x10$^{-20}$ |
| Population density: current residence | 104222 | 5.74x10$^{-05}$ | 3.28x10$^{-11}$ | 8.84x10$^{-07}$ | 8.65x10$^{-06}$ |
| Self-reported substance abuse | 633.75$^{+}$ | 0.00168$^{\alpha}$ | 3.78x10$^{-09}$ | 8.75x10$^{-09}$ | 1.34x10$^{-08}$ |
| Ever smoked cannabis | 13.04 | 0.00033 | 7.01x10$^{-08}$ | 3.02x10$^{-07}$ | 1.17x10$^{-06}$ |
| Distance travelled: birth to current | -9246200 | 9.32x10$^{-06}$ | 0.0005 | 0.0008 | 0.0002 |
| BP device ID | -670154 | 0.00014 | 0.0029 | 0.0029 | 0.0029 |
| Left hand grip strength | -888.61 | 1.15x10$^{-05}$ | 0.0054 | 0.0421 | 0.0233 |
| Population density difference: birth to current | 43655.70 | 2.11x10$^{-05}$ | 0.0061 | 0.0500 | 0.1656 |
| Breast fed | 10.41$^{+}$ | 2.08x10$^{-05\alpha}$ | 0.0585 | 0.0432 | 0.0784 |
| Population density: birth place | 91061.90 | 1.03x10$^{-05}$ | 0.0673 | 0.1019 | 0.0609 |
| Month attended baseline assessment | 33.70 | 8.32x10$^{-06}$ | 0.0989 | 0.0885 | 0.0987 |
| Leg pain on walking | 113.19$^{+}$ | 2.40x10$^{-05\alpha}$ | 0.2134 | 0.3905 | 0.2845 |
| Birth weight known | 59.44$^{+}$ | 2.88x10$^{-06\alpha}$ | 0.3291 | 0.4623 | 0.3974 |
| Birth weight | -4.56 | 3.74x10$^{-06}$ | 0.4134 | 0.4639 | 0.4058 |

+ Odds ratio for binary outcome variables; α Pseudo $R^2$ for binary outcome variables; TDEP = Townsend deprivation index; EDU = Educational attainment

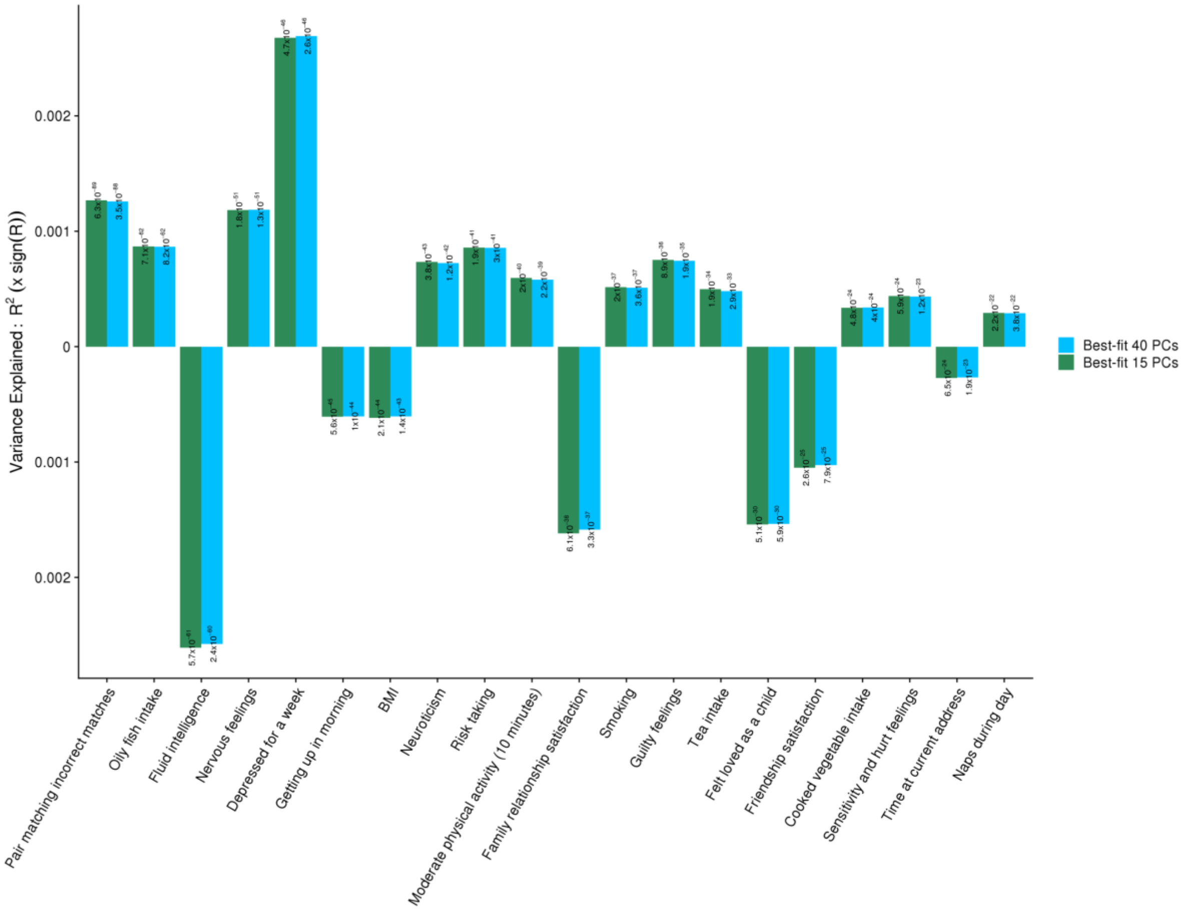


**Supplementary Figure 1.** A comparison between the results corresponding to **Fig. 1** adjusting for 15 principal components (PCs), in green, and adjusting for 40 PCs, in blue.

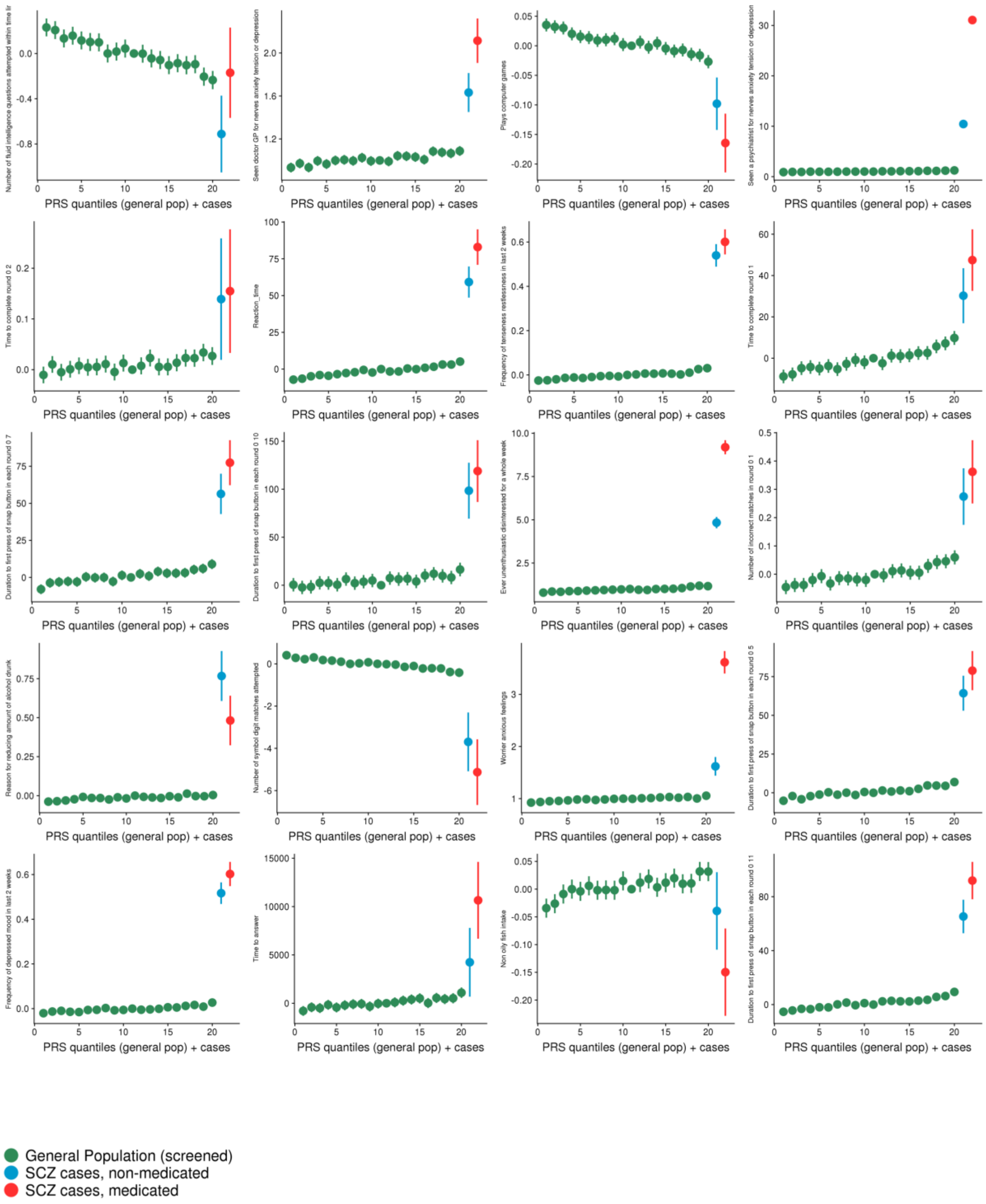


**Supplementary Figure 2.** PRS-by-trait quantile plots (in green) indicating the trends of association between the best-fit SCZ PRS and the target behavioural traits among all significant associations ($P < 1x10^{-10}$), ordered by significance, excluding those of **Fig. 1** (see **Supp. Table 1** for details). Each quantile plot is ordered from low to high genetic risk for schizophrenia, according to PRS in the target (UK Biobank) data, with each green point representing the average target trait value of a 5% quantile of the sample. Non-medicated (blue) and medicated (red), at baseline, diagnosed individuals are appended to the right end of each plot, reflecting the expected higher genetic burden of diagnosed individuals compared to unaffected individuals (see Main Text). Vertical lines represent 95% confidence intervals; these appear absent for some traits with a large range and are larger in the two categories of cases due to their smaller sample sizes.

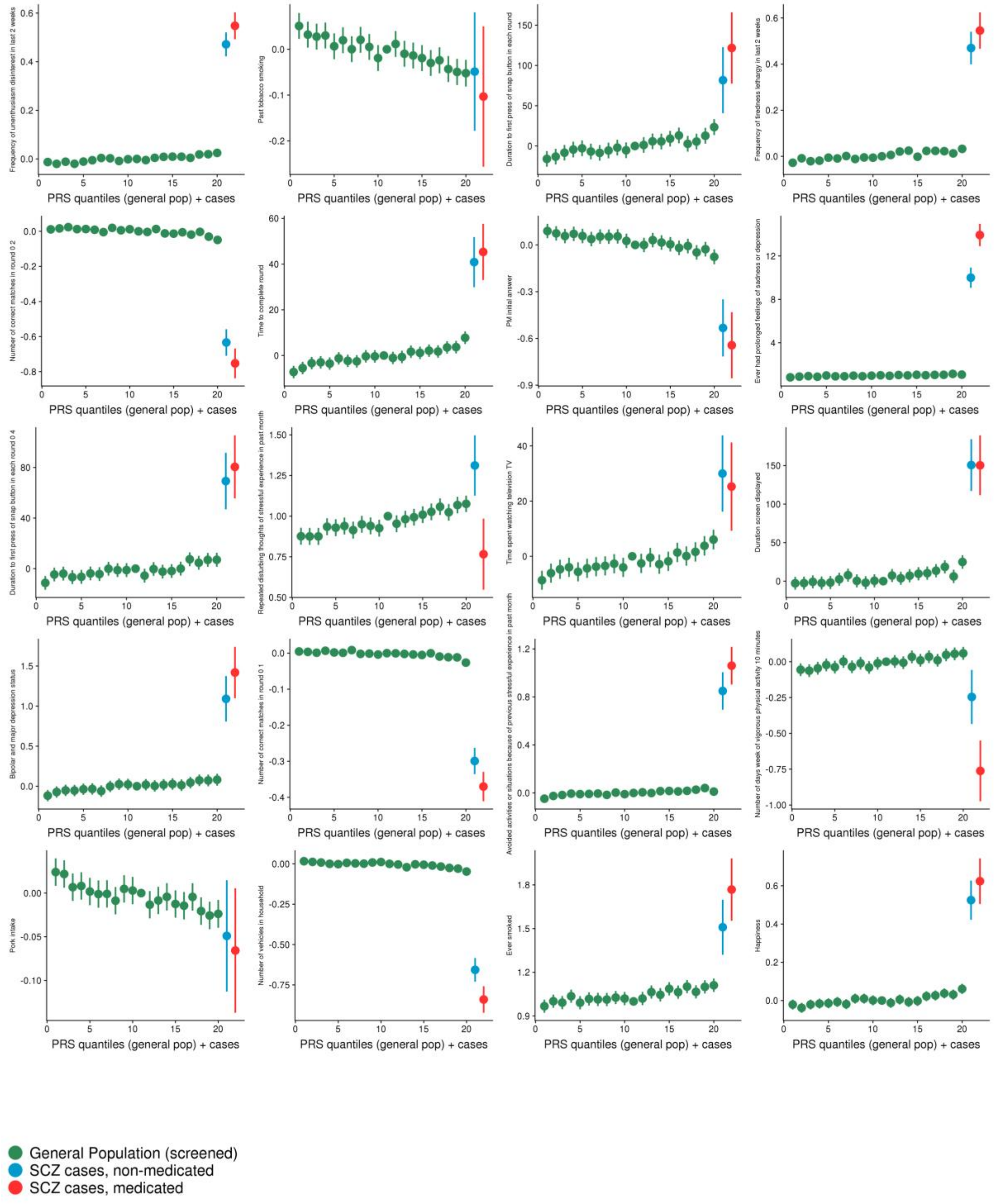


**Supplementary Figure 3.** PRS-by-trait quantile plots (in green) indicating the trends of association between the best-fit SCZ PRS and the target behavioural traits among all significant associations ($P < 1x10^{-10}$), ordered by significance, excluding those of **Fig. 1** (see **Supp. Table 1** for details). Each quantile plot is ordered from low to high genetic risk for schizophrenia, according to PRS in the target (UK Biobank) data, with each green point representing the average target trait value of a 5% quantile of the sample. Non-medicated (blue) and medicated (red), at baseline, diagnosed individuals are appended to the right end of each plot, reflecting the expected higher genetic burden of diagnosed individuals compared to unaffected individuals (see Main Text). Vertical lines represent 95% confidence intervals; these appear absent for some traits with a large range and are larger in the two categories of cases due to their smaller sample sizes.

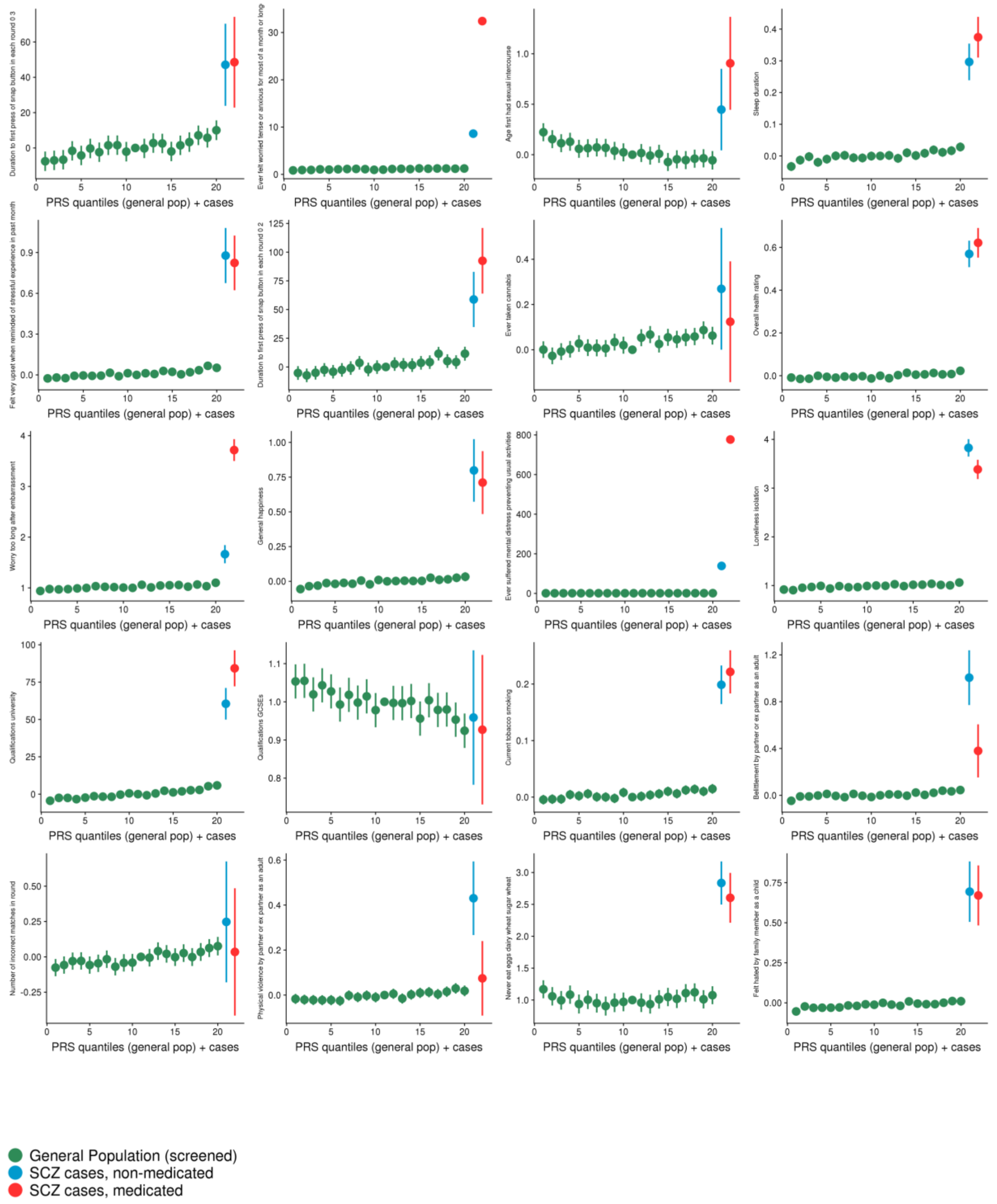


**Supplementary Figure 4.** PRS-by-trait quantile plots (in green) indicating the trends of association between the best-fit SCZ PRS and the target behavioural traits among all significant associations ($P < 1x10^{-10}$), ordered by significance, excluding those of **Fig. 1** (see **Supp. Table 1** for details). Each quantile plot is ordered from low to high genetic risk for schizophrenia, according to PRS in the target (UK Biobank) data, with each green point representing the average target trait value of a 5% quantile of the sample. Non-medicated (blue) and medicated (red), at baseline, diagnosed individuals are appended to the right end of each plot, reflecting the expected higher genetic burden of diagnosed individuals compared to unaffected individuals (see Main Text). Vertical lines represent 95% confidence intervals; these appear absent for some traits with a large range and are larger in the two categories of cases due to their smaller sample sizes.

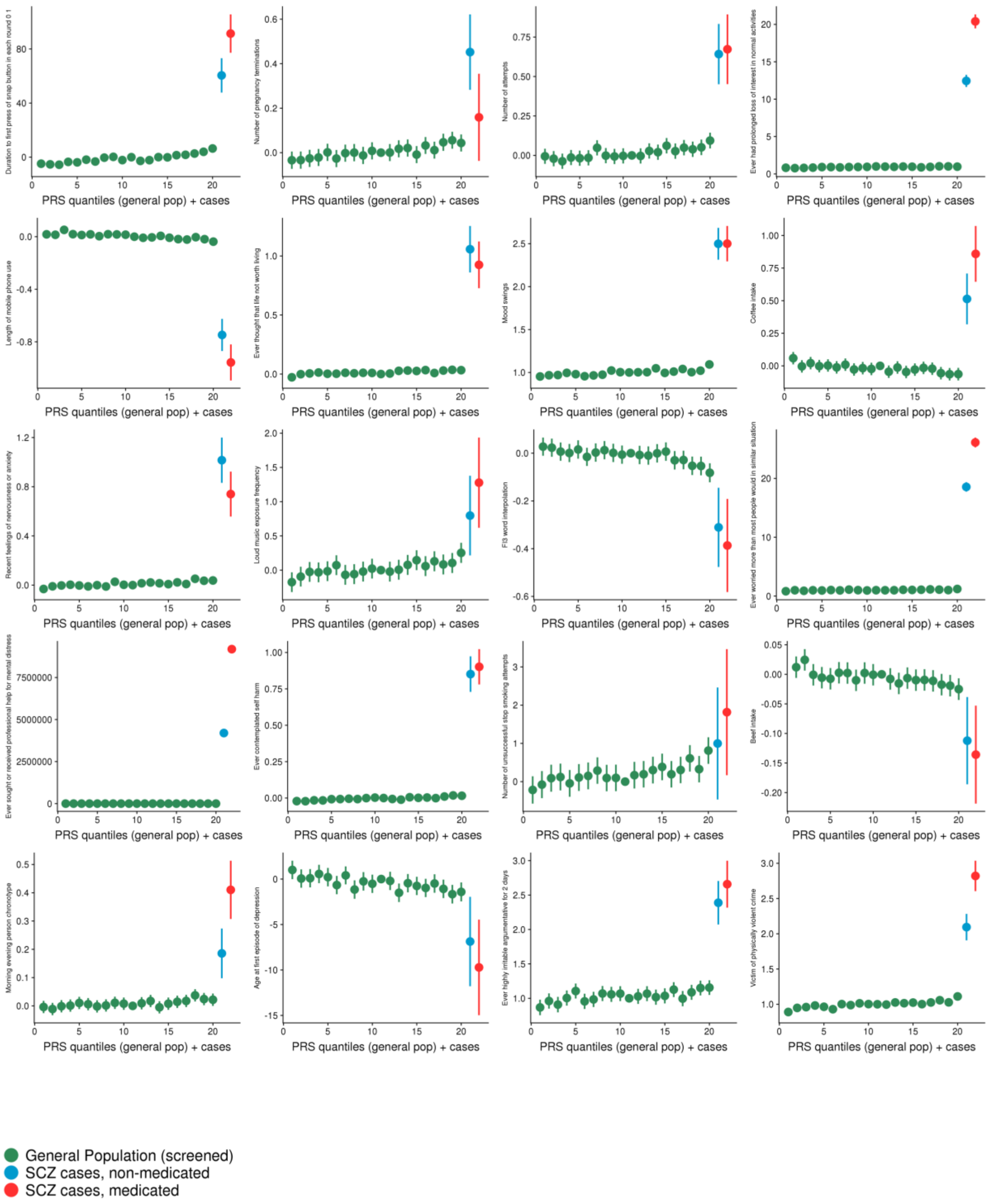


**Supplementary Figure 5.** PRS-by-trait quantile plots (in green) indicating the trends of association between the best-fit SCZ PRS and the target behavioural traits among all significant associations ($P < 1\text{x}10^{-10}$), ordered by significance, excluding those of **Fig. 1** (see **Supp. Table 1** for details). Each quantile plot is ordered from low to high genetic risk for schizophrenia, according to PRS in the target (UK Biobank) data, with each green point representing the average target trait value of a 5% quantile of the sample. Non-medicated (blue) and medicated (red), at baseline, diagnosed individuals are appended to the right end of each plot, reflecting the expected higher genetic burden of diagnosed individuals compared to unaffected individuals (see Main Text). Vertical lines represent 95% confidence intervals; these appear absent for some traits with a large range and are larger in the two categories of cases due to their smaller sample sizes.
